# Genome Functional Annotation across Species using Deep Convolutional Neural Networks

**DOI:** 10.1101/330308

**Authors:** Ghazaleh Khodabandelou, Etienne Routhier, Julien Mozziconacci

## Abstract

Deep neural network application is today a skyrocketing field in many disciplinary domains. In genomics the development of deep neural networks is expected to revolutionize current practice. Several approaches relying on convolutional neural networks have been developed to associate short genomic sequences with a functional role such as promoters, enhancers or protein binding sites along genomes. These approaches rely on the generation of sequences batches with known annotations for learning purpose. While they show good performance to predict annotations from a test subset of these batches, they usually perform poorly when applied genome-wide.

In this study, we address this issue and propose an optimal strategy to train convolutional neural networks for this specific application. We use as a case study transcription start sites and show that a model trained on one organism can be used to predict transcription start sites in a different specie. This cross-species application of convolutional neural networks trained with genomic sequence data provides a new technique to annotate any genome from previously existing annotations in related species. It also provides a way to determine whether the sequence patterns recognized by chromatin associated proteins in different species are conserved or not.

## 1 Introduction

With the advent of DNA sequencing techniques there is an explosion in availability of fully sequenced genomes. One of the major goals in the field is to interpret these DNA sequences, which is to associate a potential function with sequence motifs located at different positions along the genome. In the human genome for instance, while some DNA sequences are simply used to encode the protein sequences, most of the sequences do not code for any protein. Many of these sequences are nevertheless conserved and necessary for the correct regulation of genes. Deciphering these non-coding sequences function is a challenging task which has been increasingly achieved with the surge of next generation sequencing^1^. The 3.2 Billion base pair (bp) long human genome is now annotated with many functional and bio-chemical cues. Interpreting these annotations and their relevance in embryonic development and medicine is a current challenge in human health. While these annotations are becoming more numerous and precise, they cannot be determined experimentally for every organisms, individual and cell type. Computational methods are therefore widely used to extract sequence information from known annotations and extrapolate the results to different genomes and/or conditions. Since a few years, supervised machine learning algorithms, and notably among those deep neural networks^2^ have become a thriving field of research in genomics^3,4^. Deep Convolution Neural Networks (CNN) are indeed very efficient at detecting complex sequence patterns since they rely on the optimisation of convolution filters that can be directly matched to DNA motifs^5^. Pioneering studies illustrate the ability of CNN to reliably interpret genomic sequences^6–10^. In these studies, the training datasets are obtained in a similar fashion. For instance, Min et al.^6^ used a CNN to detect enhancers, specific sequences that regulate gene expression at a distance. Predictions were made on 1/10^*th*^ of the total number of sequences and the method reached very good scores ranking it better than previous state-of-the-art (i.e., support vector machine methods). Similar training datasets with balanced data (i.e., with a similar amount of positive and negative examples) were used in other approaches aiming at identifying promoters^7^ or detecting splicing sites^11^. While these approaches are very competitive when applied on test sets derived from the training sequences, we show here that they tend to perform poorly when applied on full chromosomes or genome-wide, which is required for the task of genome annotation.

Alternative approaches^8 10^ used unbalanced datasets for training (i.e., with more negative than positive examples) to predict DNA/RNA-binding sites for proteins and genome accessibility. In these two studies, however, the prediction performance of the model is also assessed on test sets derived from training sets, not on full genome sequences. The task of genome-wide prediction has been assessed in more recent studies aiming at identifying regulatory elements^12^ and alternative splicing sites^13^.

The methodology proposed here is inspired from these latest study and involves two important ingredients aiming at developing a tools that can achieve genome-wide predictions. Firstly, we do not take into account prediction scores obtained from test sets initially retrieved from the training sets as a quality measure. We rather assess the ability of our model to annotate a full chromosome sequence by designing a specific metric. Secondly, we optimise the ratio between positive and negative examples in order to obtain the highest prediction scores and show that this balancing optimisation is important in the case of discrete annotation features.

We then propose a new application of the method. We first train a model on a dataset corresponding to a given organism a use it to predict the annotation on the genome of a related organism, opening new opportunities for the task of *de-novo* genome annotation. As a proof of principle, we use here transcription start sites (TSS) as features. The DNA motifs around TSS can be used by proteins as recognition sites providing specific regulation of gene expression as well as precise location of the initiation of transcription^14^. The regions surrounding TSS therefore contain the information that could in principle be used to identify the TSS locations *in silico*. We show that a CNN trained on human TSS containing regions is able to recover regions containing TSS in the mouse genome and vice versa. We finally assess the generalisation of the approach to more distant species, taking as examples a fish and a bird.

## 2 Results

### 2.1 Training models for genome annotation

The problem of detecting human TSS using deep neural networks has been tackled in^7^. We first follow a similar approach, that is using a balanced dataset (see Supplementary Methods and Results for details). Briefly, the model is trained and validated on an equal number of 299 bp long positively/negatively labelled input sequences and is evaluated on a test set composed of 15% of the input data (see 4.1). Our method reaches scores of AUROC = 0.984, AUPRC = 0.988 and MCC = 0.88 which are similar to the results of^7^ (See Supplementary Methods or^15^ for the definition of these measures). In order to assess how this model would perform as a practical tool for detecting TSS on a genome-wide scale, we apply it on all the sequences along chromosome 21 (which has been withdrawn from the training set) using a 299 bp sliding window. Figure 1a illustrates the predictions of the CNN model over a typical region of 300 kbp containing 7 out of the 480 TSS of chromosome 21. Although the predictions present higher scores over TSS positions, they also present high scores over many non-TSS positions. This phenomenon makes it difficult for a predictive model to differentiate between true and false TSS. Indeed, a predictive model trained on balanced data and applied on unbalanced data tends to misclassify the majority class, hence the class with more instances in the test set. This is due to the fact that the reality is biased in the training phase during which the CNN model learns an equal number examples from the positive and the negative classes. This leads to the emergence of many false positives in a context in which the ratio between the positive and the negative class is very different (here 1:1 instead of 1:400)^16^. Facing extreme unbalances in new examples such as chromosome 21, the model fails to generalise inductive rules over the examples.

**Figure 1.**
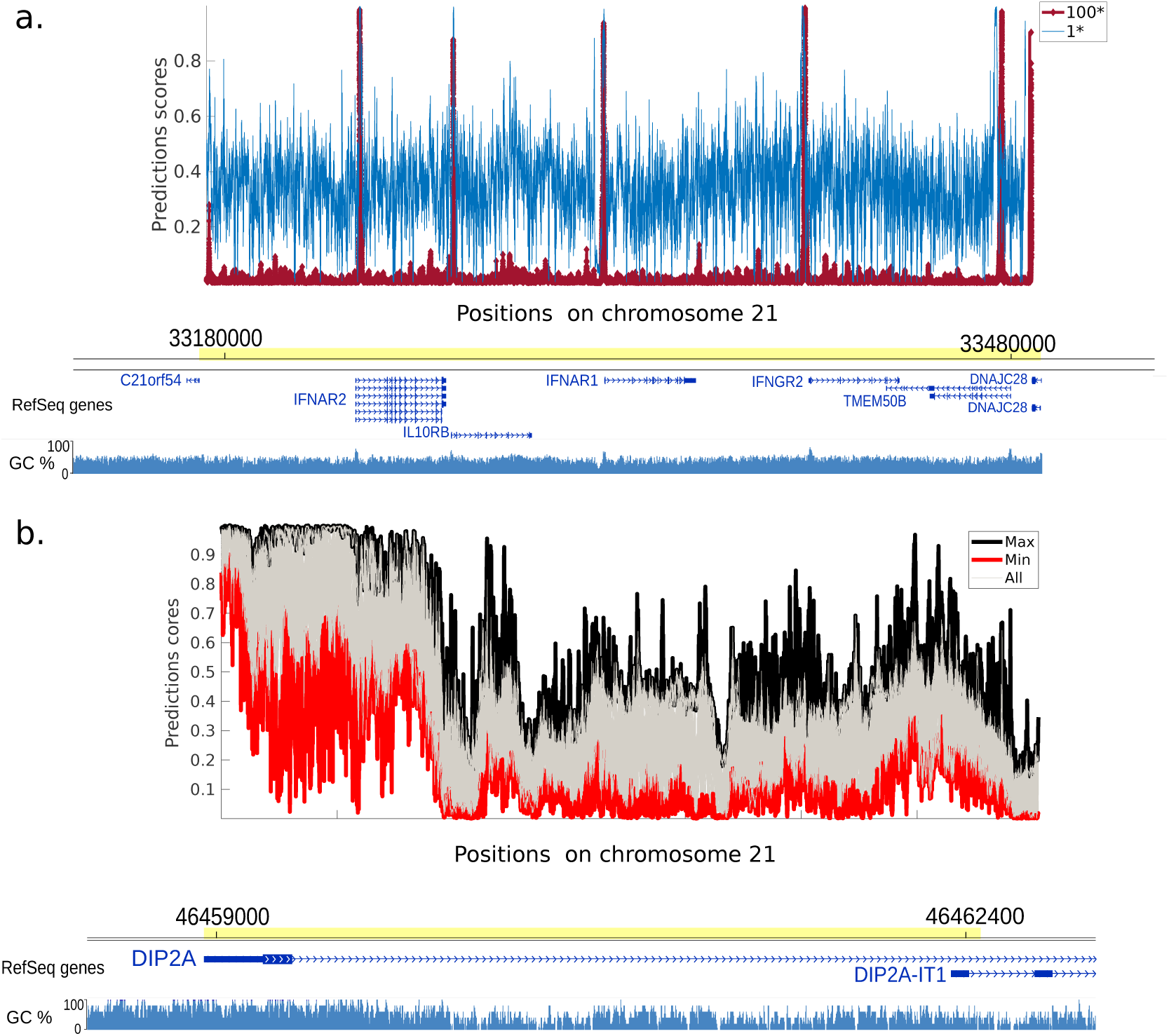
Trained CNN model prediction two regions of chromosome 21. (a) Prediction scores for balanced model 1* and unbalanced model 100*, respectively in blue and red on a 300 kbp region. The position of genes is indicated below. The GC content is indicated as blue bars below the genes. Both models detect effectively seven TSS positions, still the model 1* returns a noisier prediction. Adding more negative examples into the balanced data using the model 100* mitigates the noise while preserving the high scores over TSS. (b) Application of 30 trained CNN models over a 3.2 kbp region of chromosome 21. At each site, the maximum and minimum prediction scores are respectively displayed in black and red. Other prediction scores are plotted in grey.

To address this issue and train a network for genome annotation, we propose a heuristic approach. This approach consists in adding more negative examples into the balanced dataset to alleviating the importance of positive class in training phase and allocating more weight to the negative class. We call such datasets limited unbalanced datasets. This method, detailed in 4.1, is expected to mitigate the impact of the extreme unbalanced data on learning process. We call *Q* the ratio between negative and positive training examples and denote as *Q*^*^models trained with the corresponding ratio. For instance, on Figure 1a the model trained on the balanced data yielding to blue signal predictions is denoted as 1^*^. We train our CNN model on a 100* dataset and assess the efficiency of the trained model. As depicted on Figure 1a by a red signal, the predictions for this model display a much higher signal to noise ratio, with significant peaks over each of the 7 TSS (C21orf54, IFNAR2, IL10RB, IFNAR1, IFNGR2, TMEM50B, DNAJC28) and a much weaker signal between these sites. Predicting TSS using the 100* model is thus much efficient, since it does not generate false positive signal.

The performances of the all models evaluated using conventional metrics can be found in Supplementary Results.

### 2.2 Investigating the effect of random selection of the negative examples on predictions

Since the negative examples of the training set are randomly picked out of the genome, the performance of the model in different regions of chromosome 21 can vary for different training examples. To investigate this variation, we setup 30 balanced 1^*^ datasets and train 30 CNN separately. The 30 models are then applied over human chromosome 21 to study the fluctuations of the predictions. The variation of 30 predictions is depicted in Figure 1b. The maximum and minimum predictions for each genomic site are highlighted by black and red colours, respectively. The other 28 predictions are coloured in Gray. The first observation is that almost all predictions peak over the DIP2A TSS. However, the large gap between the minimum and maximum predictions underlines the variability of predictions obtained with different balanced datasets. This variability illustrates the uncertainty of the predictions obtained from a single CNN trained on a balanced dataset. This highlights the need to use limited unbalanced datasets for the task of genome annotation.

### 2.3 Comparing 1* and 100* models over a full chromosome

Models trained on 1* and 100* sets are applied over the full chromosome 21 and the predictions around TSS are presented as heat-maps. In Figure 2a,b each horizontal line corresponds to the standard score of the predictions computed over ± 5000 bp of each TSS of chromosome 21 for the models 1* and 100*, respectively. The central axis indicates the exact position of the TSS and positions around the axis indicate the neighbouring regions. While the model 1* Figure 2a presents a noisier signal around TSS positions, the model 100* Figure 2b displays a higher signal to noise ratio. To investigate the performance of different models on a genome-wide scale we devised a custom metric *λ* which measures the average signal to noise ratio increase around TSS (see Methods for the definition of *λ*).

**Figure 2.**
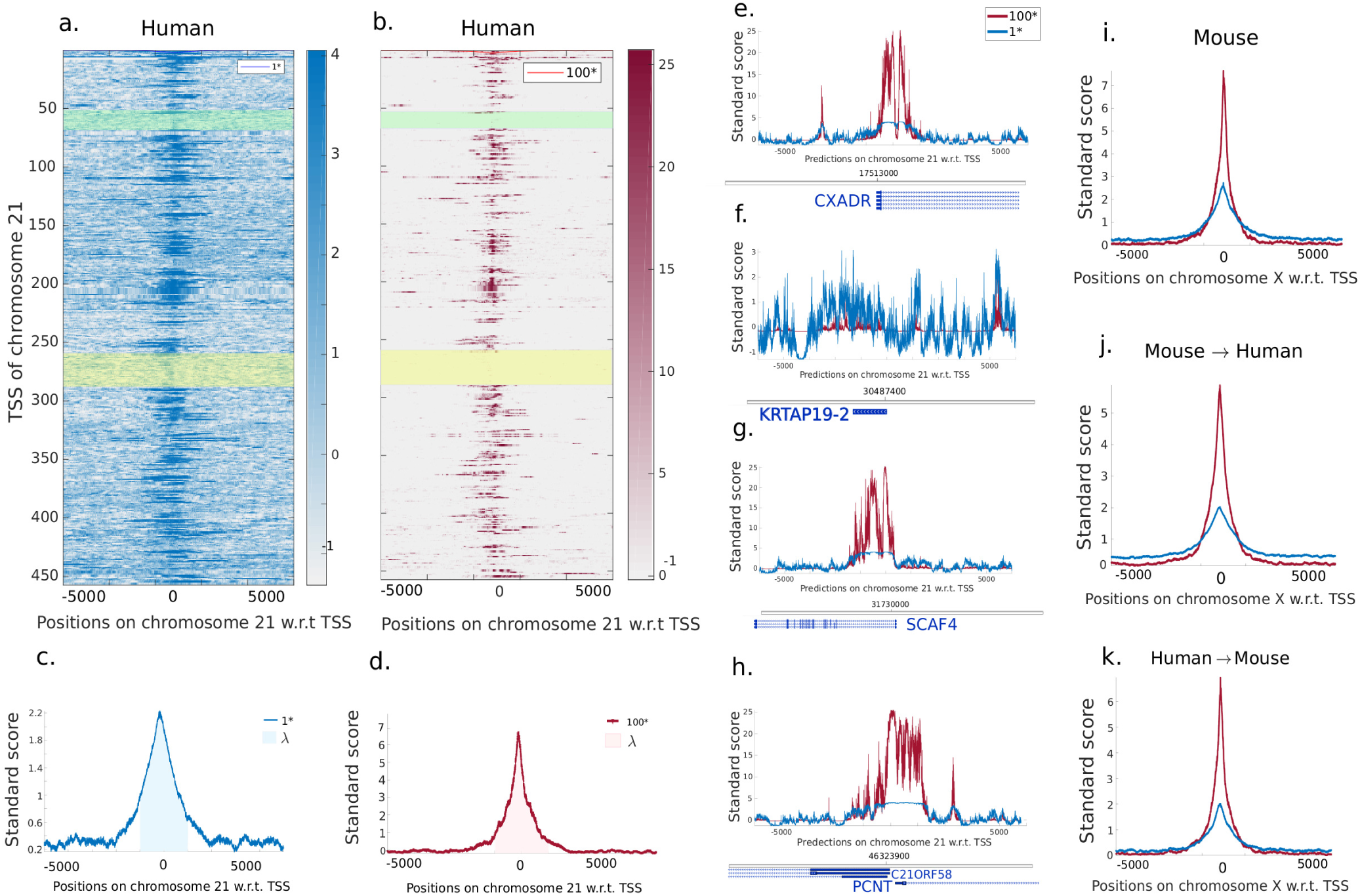
Comparison of the 1* and 100* models predictions over chromosome 21. (a) and (b) Heat maps depict the standard score of the prediction for respectively the 1* and 100* models on 5000 bp flanking each TSS of chromosome 21. (c) and (d) Averaged standard score of the predictions over each TSS of chromosome 21. (e-h) Zoom on regions around selected TSS. Genes are indicated at the bottom of each plot. (i-k) Averaged standard score of the predictions over each TSS of mouse chromosome X (i) and for networks trained on mouse/human chromosomes (except X) and applied on human/mouse chromosome X (j,k).

Figure 2c,d illustrate the average of the standard score over all the TSS of chromosome 21 for the model 1* and 100*, respectively and *λ* denotes the average of this average over a r=2 kbp region centred on the TSS. A larger *λ* score corresponds to a better signal to noise ratio. In this particular case, we find a *λ* score of 1.49 and 2.99 for the 1* and 100* model, respectively.

To illustrate the variability of prediction scores achieved around different TSS, we choose four representatives TSS within the chromosome. The first TSS corresponds to the gene CXADR, shown in Figure 2e. While the prediction of model 1* results in a poor averaged standard scores over all positions, the averaged standard score of model 100* strongly peaks around the TSS position and shows low variations over non-TSS positions. Figure 2f depicts the second selected TSS corresponding to the KRTAP19-2 gene. This gene is part of a cluster of similar genes belonging to the family of Keratin Associated Proteins (highlighted by a yellow rectangle on Figure 2a,b). For this particular cluster, the predictions are poor for both 1* and 100*, probably reflecting a specific TSS signature that has not been grasped by the model. Another example of gene cluster with a poor prediction score for TSS is the t-RNA cluster, highlighted in green. Figure 2g,h displays the predictions around the TSS of the SCAF4 and, PCNT and C21ORF58 genes, respectively. On these more typical TSS the 100* model shows a higher signal to noise ratio than the 1* and regions containing TSS are faithfully detected. These regions often stretch over 1 kb while our training sequence centered on each TSS is only 299bp long. This could indicate the presence either of alternative TSS close to the annotated TSS or of similar sequence patterns in broader regions surrounding the TSS.

### 2.4 Learning and predicting in human and mouse

To show the potential of our annotation method in a different context, we replicate a similar TSS analysis in mouse. Models with values of *Q* ranging from 1 to 100 trained on mouse chromosomes (except X) are applied over the mouse chromosome X to assess the model performance (see Figure 2i, Supplementary Figure 1 and Supplementary Figure 2a,d,g). The averaged standard score *λ* reaches values of 1.47 and 2.18 respectively for the 1* and 100* models in quantitative agreement with the model performance in human. We next determine the possibility of predicting TSS in one organism with a network trained on a different albeit related organisms. To this end, the mouse trained model is applied on human chromosome X and the human trained model is applied on mouse chromosome X. The two chromosomes carry homologous genes, the number of annotated TSS varies with a total of 4,968 TSS in human and 2,005 TSS in mouse. While the model trained and applied on mouse shows a better signal to noise ratio, the same model applied to human chromosome X still captures most of the TSS and gives a *λ* score of 2.28 for the 100* model (see Figure 2j and Supplementary Figure 2b,e,h). Similarly, the models trained on human capture most of TSS on the mouse X chromosome as shown in Figure 2k and Supplementary Figure 2c,f,i and reaches a *λ* score of 2.04 for the 100* model. In all cases, the signal to noise ratio is improved in the 100* models. The values of *λ* for the cross-species comparison on chromosome X are summed up in Supplementary Figure 1. The human model applied on human provides the highest scores for both 1* and 100* models probably a signature of an overall better TSS annotation. In all cases, *λ* gradually increases for 10* up to 100* datasets. It should be pointed out that since the performance of models 30* and 100* varies slightly, the 30* model can be used instead of 100* to perform cost-effective computations.

### 2.5 Evaluation of the prediction for different TSS classes

The potential of our trained networks to recover TSS containing regions along the human and mouse genome is assessed in the previous parts without any distinction between different TSS classes. Since we find that some TSS are better predicted than others (Figure 1), we compute the *λ* score independently for the two main classes of TSS: mRNA-TSS and ncRNA-TSS. While *λ* is higher for the mRNA-TSS class, the model is versatile and is also able to predict the ncRNA-TSS (Figure 3b). In human and mouse, mRNA-TSS are found in different classes, that can be derived from the CG di-nucleotide (CpG) content of the region flanking the TSS. High CpG regions, also called “island” can be methylated and play an important role in gene regulation^17^. Figure 3a displays the distribution of CpG number in 299 bp regions surrounding the all mRNA-TSS for the mouse and human X chromosome. From this distribution, we identify three classes of mRNA-TSS with respectively a high, medium and low CpG content. High CpG TSS correspond to genes regulated by DNA methylation and have been shown to exhibit a different pattern of chromatin modifications^18^. Assessing the performance of the model for three different classes, we find that stronger scores are obtained for CpG richer TSS (Figure 3b). The worst performing TSS are low CpG content TSS which are not recovered at all by our model. This points at a larger diversity and complexity of motifs in low CpG content regions surrounding TSS. It would be interesting to test if a larger kernel size of the convolution filter would allow for a better detection of these regions.

**Figure 3.**
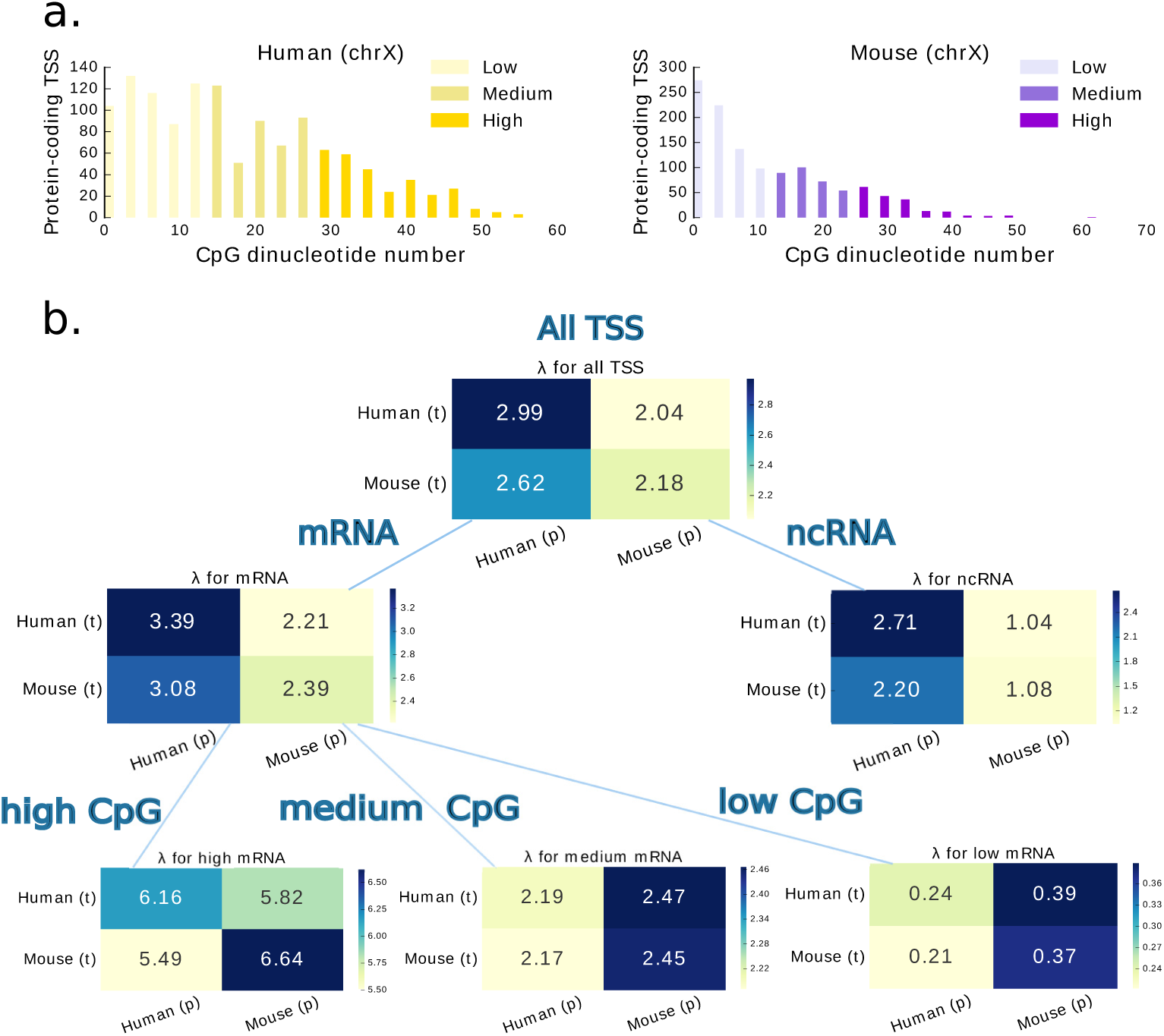
Evaluation of the model performances for different RNA classes. (a) CG di-nucleotide (CpG) number in 299bp regions centred on mRNA-TSS in test chromosomes (X). Three populations (low, medium and high) can be identified. (b) Each box represent the *λ* values obtained when training the network on the organism corresponding to each line and predicting the TSS on the chromosome X of the organism corresponding to the column; *λ* scores are also computed for mRNA- and ncRNA-TSS separately and finally for mRNA-TSS divided into three classes based on the CpG density populations identified in (a).

### 2.6 Application of the approach to other vertebrates

The performances of a CNN trained on human TSS to recover mouse TSS is not surprising given the similarity between the two mammalian genomes^19^. We next set out to apply the same methodology on more diverse species, including a bird and a fish (Figure 4). Four CNN are trained on all the TSS of the Human, Mouse, Chicken and Danio rerio (Zebrafish) genomes which provide the most comprehensive TSS annotations for mammals, birds and fishes. These four models are then applied genome wide on each of the four species and the *λ* metric is computed for each chromosomes independently, using a *r* value of 400 bp (see Methods).

**Figure 4.**
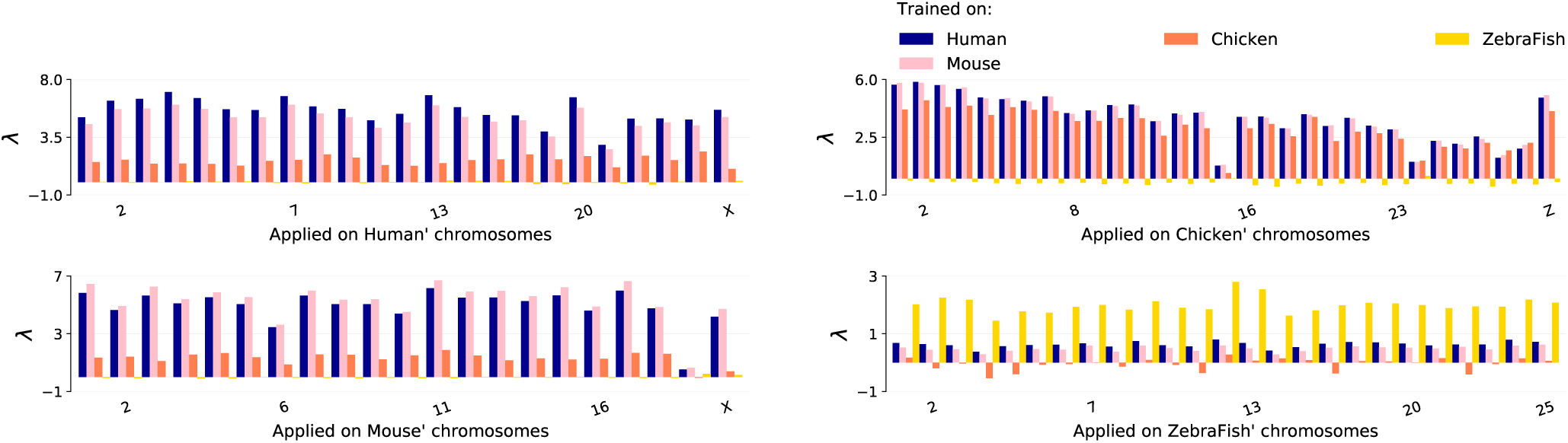
The CNN model trained on four different species: Human, Mouse, Chicken and Zebrafish is then applied over each chromosomes of these four species. In this figure, the size of the window *r* on which *λ* is computed is equal to 400 bp.

The results for the human and mouse genomes are very alike, with only a slightly better performance when the model trained on a species is applied on the same specie. The model trained on the Chicken genome performs less well when applied on the mammalian genomes and the model trained on the Zebrafish genome is not able recover the mammalian TSS with a *λ* value of 0.

When applied on the Chicken genome, the mouse and human models surprisingly outperform the chicken model, probably because the TSS annotation is better in the two mammals so that the training phase is more efficient. This result highlights the potential of the method when used across different species when one specie’s genome is more precisely annotated.

When applied on the Zebrafish genome on the other hand, the human, mouse and chicken models all show poor performances while the Zebrafish model does a pretty good job. This is in line with the fact that the CpG composition of Zebrafish regions around TSS if very different than in birds and mammals. CpG islands, which are high density CpG regions, are found upstream many TSS for coding genes in birds and mammals while they are absent in fishes. All together, these results suggest that the molecular machinery that is able to interpret the genome sequence in order to find the start sites of genes has a similar activity in human, mouse and chicken but in a different activity in fishes.

## 3 Discussion

With the surge of DNA sequencing technologies, one million of genomes are now available and millions of gigabases are sequenced every year to annotate these genomes with functional marks^20^. It has not escape the notice of many computational biologists that deep neural networks is a key tool to deal with this exponentially increasing amount of data^20^. One of the practical issues when applying CNN on genomic sequences is the unbalanced data, a well-known issue in the machine learning literature^16, 21, 22^. In the present paper, we address this problem using TSS as a case study. Indeed, the TSS occupy only a few locations on the genome (31,037 TSS for human) leading to extreme unbalances in dataset (i.e., 1/400 is the ratio of TSS-containing 299 bp windows to non-TSS in the human genome). This issue is usually referred to as a rare events or rare samples. In this scenario, the lack of examples of the minority class (i.e., true TSS) deteriorates the learning process as conventional machine learning algorithms usually measure the model performance on the most representative class (i.e., non-TSS) leading to biased or inaccurate prediction of the minority class. To deal with this disparity, we adopt a sub-sampling strategy to decrease the frequency of the majority class samples (non-TSS) improving thereby identification of the rare minority class samples (TSS). We show that this method achieves very good performances for CNN models in predicting TSS and non-TSS sequences genome-wide without requiring knowledge of any specific TSS features. Since the convolution filters are able to automatically capture sequence motifs and other significant characteristics of genomic sequences, this approach can be easily extended to identify other functional regions in any annotated genomes.

TSS (or any other regulatory elements in the genome) are recognised by the cellular machinery enabling the transcription of genes to start at precise locations on genome. These locations found by macromolecular complexes, which contains proteins able to recognise specific DNA motifs. We show that our method can be efficiently used across different genomes, that is training the model in one genome and applying it to another genome. We use human and mouse TSS as case study and first apply both models on chromosome X of each organism. While the sequence of this chromosome has evolved differently in both species, many genes are homologous. The fact that we are able to recover TSS in mouse/human with a model trained in the other organism, suggest that the machinery capable of recognising TSS in both organism is overall conserved. We also show that this methodology can be apply to more distant species, and use as examples a bird and a fish. Our results point toward a high similarity between mammals and birds while fishes TSS cannot be predicted from mammals and bird sequences. While the genome sequence conservation can be computed directly from DNA sequences, our method provides a new tool to address the conservation of the activity of the nuclear machinery that interprets the sequences. We expect that such a tool can be widely used in the future both by evolutionary biologists and by molecular biologists interested in the evolution of chromatin associated proteins.

## 4 Methods

### 4.1 Input Generation

The TSS positions are collected from the reference genomes for human (hg38) and mouse (mm10) species. Genome sequences data are available at http://hgdownload.soe.ucsc.edu/. TSS positions over the entire human and mouse genomes data are available at http://egg.wustl.edu/, the gene annotation is taken from RefGene. As positive input sequences, we use regions of 299 bp flanking TSS (i.e., ± 149 bp around the TSS) which are supposed to contain multiple sequence signals indicating the presence of a TSS to the transcription machinery of the cell. Overall, 31,037 TSS positions are extracted on both DNA strands (15,798 for positive strand and 15,239 for negative strand). In a similar fashion, we extract 25,698 TSS positions out of the mouse genome (12,938 for positive strand and 12,760 for negative strand). In order to generate the negative class, we perform a sub-sampling strategy on both genomes to set up a balanced dataset. To do this, we select *Q* × 31, 037 299 bp long regions at random positions, insuring that these regions do not contain a TSS. The odds of getting at random a genomic region containing a TSS are close to 0.25%. For *Q* = 1, there is an equal number of negative and positive class examples. Unbalanced datasets are produced using different values of *Q* ranging from 1 to 100.

### 4.2 Convolution Neural Network (CNN)

A prediction model is implemented using CNN in order to predict the presence of a TSS in a DNA sequence of size 299 bp. Our deep convolutional neural network typically has an input shape of *c* × *b*. In our model architecture, summarised in Fig 5, the input is a matrix for which *c* = 4 is the number of different nucleotides and *b* = 299 is the length of the input sequence. The nucleotide sequences are one hot encoded so that A=(1,0,0,0), T=(0,1,0,0), G=(0,0,1,0), and C=(0,0,0,1). The training set contains *N* samples of labelled-pairs (*X* ^(*n*)^, *y*^(*n*)^), for *n* ∈ {1, …, *N*}, where *X* ^(*n*)^are matrices of size *c*(= 4) × *b*(= 299) and *y*^(*n*)^ ∈ {0, 1} Each sample in the matrices *X* ^(*n*)^ is labelled by *y*^(*n*)^ = 1 when it corresponds to a TSS containing region and *y*^(*n*)^ = 0 otherwise. The convolutional layers carry out a series of sliding windows operation (convolution) using *k* kernels each of size *s* to scan motifs all over positions *p*. The first layer kernels perform convolution on *s* successive input sequences positions *p* ∈ {1, …, (*b -s* + 1)} to recognise relevant patterns that are optimized during the training phase. The next convolutional layers scan low-level features in the previous layers in order to model high-level features. This filtering operation generates an output feature map of size *k* × (*b -s* + 1) for an input sample *X* of size *c* × *b*. The feature map ℳ = *f*_*conv*_(*X*) resulting from the convolution operation is computed as follows:

**Figure 5.**
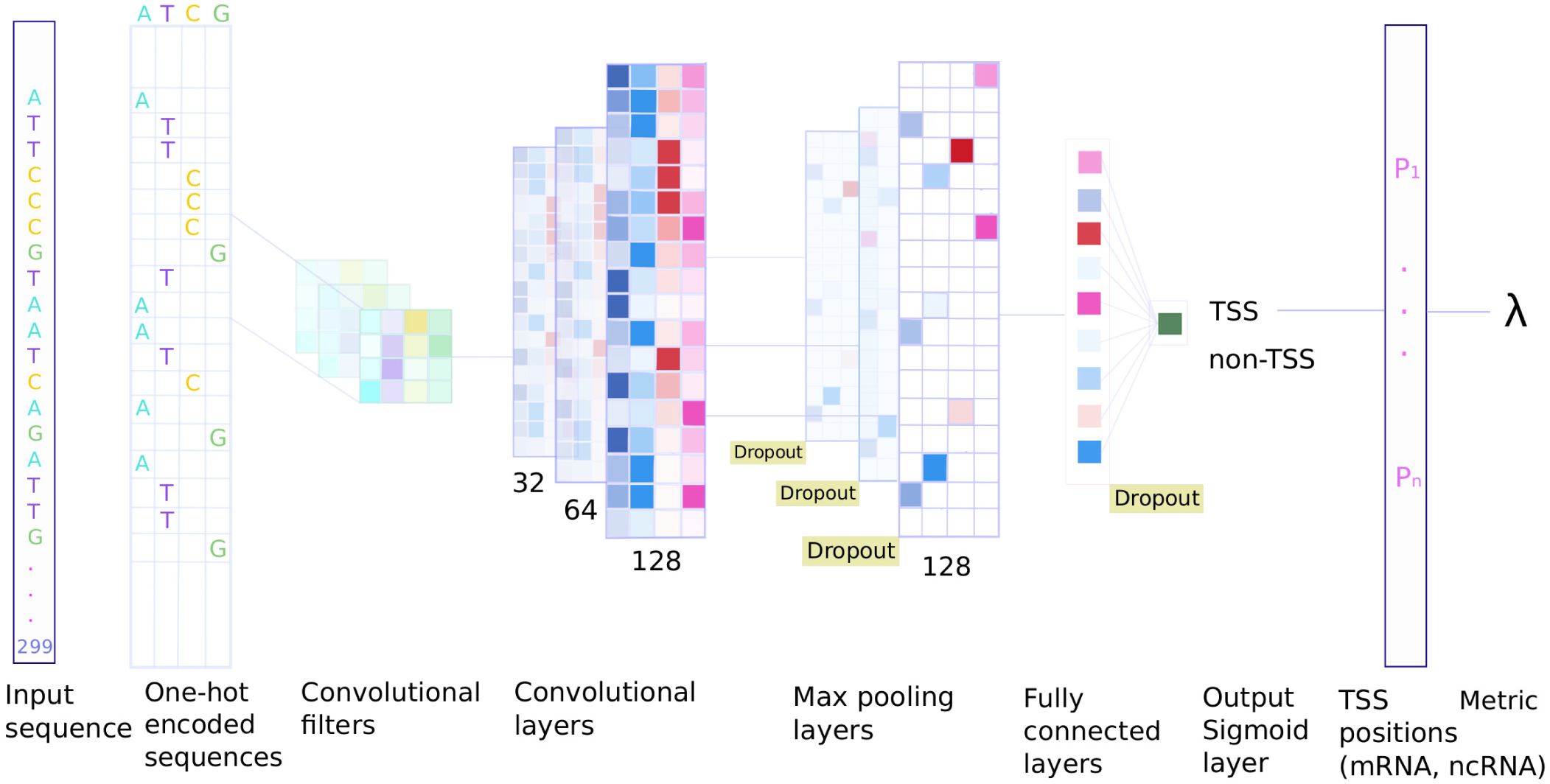
Overview of the CNN model. Input 299 bp long-sequences are one hot encoded into a 4× 299 input matrix. The first layer performs a convolution on each input matrices to recognise relevant patterns. The next convolutional layers model the interplay among these patterns to form higher-level features. Max-pooling layers reduce the dimensions of the patterns lowering computational cost and promoting high-level features detection. Dropout layers discard randomly some outputs of previous layers to avoid over-fitting. The model is optimised to correctly label input sequences as TSS or non-TSS. The output layer of the network then gives a probability for each 299 bp region to contain a TSS over the full chromosome length. The metric *λ* is computed from the results of the model to measure the average z-transformed probability around all the known TSS in order to measure the performance of the model.

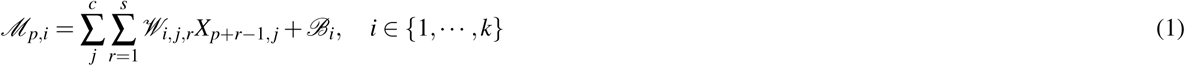

where *𝒲* denotes the network weights with size (*k* × *c* × *s*) and *ℬ* denotes the biases with size (*k* × 1) (see e.g.^2^). After each convolution layer a non-linear function is applied to the output, here a Rectified Linear Unit (ReLU). This activation function computes *f*_*ReLU*_ (*ℳ*) = *max*(0, *ℳ*) to incorporate non-linearity by transforming all negative values to zero. In order to reduce the input dimension and provide an abstract form of the representation, we apply a max-pooling process with pool size *m* over this output of *f*_*ReLU*_ (*ℳ*). Max-pooling reduces the input parameters to a lower dimension and provides basic translation invariance to the inner representation resulting in less computational cost and promotes high-level patterns detection. In our model, max-pooling layer reduces a large region into a smaller representation by taking the maximum values of the range in pooling size.

One of the important issues of any learning algorithm is over-fitting. Overfitting occurs when you achieve a good fit of your model on the training data, while it does not generalise well on new, unseen data. To deal with this issue, a regularisation procedure called dropout is usually used^23^. In the training step, some outputs of each layer are randomly masked while the remaining information is fed as the inputs to the next layer.

The final step is using a fully connected layer to learn a model to map DNA sequences *X* ^(*n*)^ onto TSS positions *y*^(*n*)^. While convolutional layers learn high-level features, the fully connected layer deals with linear and non-linear combinations of high-level features arising from the convolutional layers in order to make the final prediction. In order to learn all hidden features, our network has several hidden layers and the last layer is the output layer, which through a sigmoid function 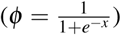 generates the class scores for all input sequences. In the training phase, the weights and biases of the convolution layers and of the fully connected layer are updated via a back-propagation process so that the loss, which measure the discrepancy between the network predictions and actual values, decreases. The loss function computes an average over the losses for every individual example. The metric of binary Cross Entropy (log loss) is commonly-used to measures the performance of a classification model which output is a probability value between 0 and 1. It is computed as:

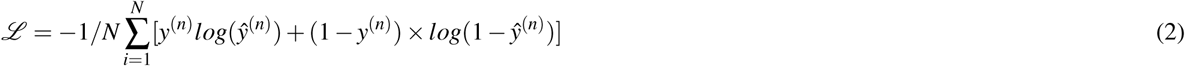

where *ŷ*^(*n*)^ is the estimated scores for the input samples *X* ^(*n*)^.

### 4.3 Implementation

We implement CNN using Keras library and Tensorflow^24^ as back-end. For a faster training a GTX 1070 Ti GPU is used. We use Adaptive Moment Estimation (Adam) to compute adaptive learning rates for each parameter^25^. Adam optimiser is an algorithm for first-order stochastic gradient-based optimisation of functions, based on adaptive estimates of lower-order moments. Our CNN architecture (see Fig5) consists of three convolutional layers of size 32, 64 and 128 kernels each of shape (4 × 4) with sliding step equals to 1. The output from convolutional is entered into a max-pooling layer with pooling size (1 × 2). After each max-pooling layer a ReLU layer is used following by a dropout with a ratio of 0.2. Finally, the output of the last pooling layer is flattened to 1D via a Flatten layer and passed through a fully connected (Dense) layer with 128 neurons. The output layer contains a single neuron. It uses a sigmoid activation function to make predictions by producing a probability output of being TSS or non-TSS. The network architecture is detailed in Supplementary Table 1. The models are trained for 150 epochs and they mostly converge early (around 30-35 epochs). All the data and the program are available at https://github.com/StudyTSS/DeepTSS/.

### 4.4 Genome wide performance measure

Different measures have been developed in order to assess the performance models applied on conventional test sets, i.e. tests sets derived from a subset of the initial data (see Supplementary Methods). In our case, the test set is different from the training set in the sense that we apply our model on all the 299 bp windows spanning full unseen chromosomes and eventually chromosomes from other species. We therefore developed a measure to evaluate the performance of the trained models in this case. This custom metric, called *λ*, measures the enhancement of the predicted signal specifically in a 2 kbp region surrounding the known TSS. To compute *λ*, we first compute the genome-wide standard score 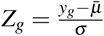 from the predictions *y*_*g*_ where *g* denotes positions on the genome, and 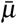 and *σ* stand for the predictions mean and standard deviation, respectively. We extract *𝒵*_*TSS*_, the *Z*_*g*_ signal over 10 kbp windows centred on each TSS of the test region, e.g. a full unseen chromosome. We average element wise *𝒵*_*TSS*_ over all TSS. This gives us *S*, the average of the Z-transformed prediction score in a 10 kb window around TSS. In order to measure the signal increase close to the TSS, that we call *λ*, we compute the average of the curve *S* on a region of *r* = 2 kbp centred on the TSS. A higher value of *λ* corresponds to a higher signal to noise ratio in a 2 kbp region around the TSS.

## Acknowledgements

We would like to thank Léopold Carron for helping us with datasets, Hugues Roest Croeluis for discussions and Michel Quaggetto for technical support.

## Additional Information

### Author Contributions Statement

GK and JM conceived and designed the methodology. GK performed the experiments. GK, ER and JM analyzed the results. GK and JM wrote the paper. GK and JM revised the manuscript.

### Data availability statement

Genome sequences data and gene annotation are available at http://hgdownload.soe.ucsc.edu/ and http://egg.wustl.edu/.

